# IFITM3 protects the heart during influenza virus infection

**DOI:** 10.1101/518548

**Authors:** Adam D. Kenney, Temet M. McMichael, Alexander Imas, Nicholas M. Chesarino, Lizhi Zhang, Lisa E. Dorn, Qian Wu, Omar Alfaour, Foued Amari, Min Chen, Ashley Zani, Mahesh Chemudupati, Federica Accornero, Vincenzo Coppola, Murugesan V.S. Rajaram, Jacob S. Yount

**Affiliations:** Department of Microbial Infection and Immunity, The Ohio State University, Columbus, OH 43210, USA; Department of Physiology and Cell Biology, The Ohio State University, Columbus, OH 43210, USA; Genetically Engineered Mouse Modeling Core, The Ohio State University and James Comprehensive Cancer Center, Columbus, OH 43210, USA; Department of Cancer Biology and Genetics, The Ohio State University, Columbus, OH 43210, USA

**Author notes:** Correspondence to J.S.Y. or M.V.S.R.

## Abstract

Influenza virus primarily targets the lungs, but dissemination and damage to heart tissue is also known to occur in severe infections. Despite this knowledge, influenza virus-induced cardiac pathogenesis and its underlying mechanisms have been difficult to study due to a lack of small animal models. In humans, polymorphisms in the gene encoding interferon-induced transmembrane protein 3 (IFITM3), an antiviral restriction factor, are associated with susceptibility to severe influenza, but whether IFITM3 deficiencies contribute to other aspects of pathogenesis, including cardiac dysfunction, is unknown. We now show that IFITM3 deficiency in a newly generated knockout (KO) mouse model exacerbates illness and mortality following influenza A virus infection. Enhanced pathogenesis correlated with increased replication of virus in the lungs, spleens, and hearts of KO mice relative to wildtype (WT) mice. IFITM3 KO mice exhibited normal cardiac function at baseline, but developed severely aberrant electrical activity upon infection, including decreased heart rate and irregular, arrhythmic RR (interbeat) intervals. In contrast, WT mice exhibited a mild decrease in heart rate without irregularity of RR intervals. Heightened cardiac virus titers and electrical dysfunction in KO animals was accompanied by increased activation of fibrotic pathways and fibrotic lesions in the heart. Our findings reveal an essential role for IFITM3 in controlling influenza virus replication and pathogenesis in heart tissue and establish IFITM3 KO mice as a powerful model to study virus-induced cardiac dysfunction.

## Introduction

Influenza virus is among the top ten causes of human mortality, and illness caused by the virus is estimated to cost the US economy as much as $80 billion per year due to lost productivity (Molinari et al., 2007). Although influenza virus primarily infects the lungs, cardiac complications of infection are also well documented (Karjalainen et al., 1980; Kodama, 2010; Lucke et al., 1919; Oseasohn et al., 1959; Paddock et al., 2012; Ukimura et al., 2010; Ukimura et al., 2013). These cardiac pathologies may result from systemic inflammation, stress caused by infection, or direct infection of heart tissue. Indeed, influenza virus is known to be a cardiotropic virus that can disseminate from the lungs to infect heart tissue, particularly during severe infections (Fislova et al., 2009; Iwatsuki-Horimoto et al., 2018; Kobasa et al., 2007; Kotaka et al., 1990). Thus, the virus can cause myocarditis and cardiac dysfunction even in individuals without pre-existing cardiovascular disease. Signs of myocarditis have been observed in up to 13% of individuals hospitalized with influenza virus infections(Karjalainen et al., 1980; Kodama, 2010; Ukimura et al., 2013). Further, significant signs of myocarditis at autopsy has been reported for up to 48% of fatal seasonal influenza cases (Oseasohn et al., 1959; Paddock et al., 2012; Ukimura et al., 2010; Ukimura et al., 2013), and a landmark autopsy study reported severe cardiac damage in a majority of 126 patients examined after succumbing to infection with the 1918 pandemic H1N1 influenza virus (Lucke et al., 1919). Infections of cynomolgus macaques with the 1918 virus showed that the virus disseminated to the heart as early as 3 days post-infection (Kobasa et al., 2007). In sum, severe influenza virus infections are associated with heart tissue infection and pathology, but the underlying mechanisms and susceptibility factors for these effects are poorly understood.

Influenza A virus infections of extrapulmonary tissues, including the heart, spleen, kidney, thymus, and brain have been reported for various mouse models of infection (Fislova et al., 2009; Kotaka et al., 1990; Tundup et al., 2017). However, relevant models in which significant cardiac pathogenesis occurs are lacking, thus hindering advancement in the study of cardiac complications of influenza virus infection. For example, infection of mice with the pathogenic PR8 strain of influenza virus (A/Puerto Rico/8/1934 (H1N1)) at a sublethal dose results in spread of the virus to the heart and replication to low levels (Fislova et al., 2009; Kotaka et al., 1990), but the resulting myocarditis is mild and resolves quickly as virus is cleared (Kotaka et al., 1990). Infections with extreme virus doses can cause increased pathogenesis and lethality, but the rapid death that is induced by such infection regimes may not involve the same pathogenic mechanisms that are initiated during a longer course of infection with a lower dose (Vogel et al., 2014; Woods et al., 2016). Likewise, animals lacking the type I interferon (IFN) receptor or downstream signaling molecules experience increased pathogenicity upon infection (Garcia-Sastre et al., 1998; Szretter et al., 2009), but deficiencies in these animals correspond to genetic defects that are rarely seen in humans (Kenney et al., 2017). Overall, there is a need for an animal model displaying significant cardiac complications of influenza virus infection in which the susceptibility of the animal is relevant to human infections, and in which a physiologically relevant virus dose is administered via the intranasal route.

Single nucleotide polymorphisms (SNPs) in the *IFITM3* gene or its promoter are the only genetic defects that have been reproducibly associated with severe influenza in humans (Allen et al., 2017; Everitt et al., 2012; Kenney et al., 2017; Pan et al., 2017; Wang et al., 2014; Zani and Yount, 2018; Zhang et al., 2013). IFITM3 directly restricts influenza virus infection by inhibiting virus entry into cells, and it also provides a secondary function in dampening tissue-damaging inflammatory cytokine responses (Chesarino et al., 2017; Desai et al., 2014; Feeley et al., 2011; Jiang et al., 2017; Li et al., 2013; Stacey et al., 2017). The rs12252-C SNP has been suggested to cause IFITM3 mislocalization and thus an inability to inhibit influenza virus infections (Chesarino et al., 2014; Everitt et al., 2012). In the Han Chinese population, this SNP is homozygous in 20% of individuals (Zani and Yount, 2018). Another SNP, rs34481144-A, located in the IFITM3 promoter, is homozygous in 4% of people with European ancestry (Zani and Yount, 2018). These individuals produce low levels of IFITM3, making them more susceptible to infection (Allen et al., 2017). Given that numerous studies have linked IFITM3 defects to severe influenza virus infections, and given that severe influenza virus infections may include cardiac complications, we sought to investigate whether IFITM3 plays a role in protecting the heart during infection. Using our newly developed IFITM3 KO mice, we have discovered that IFITM3 limits replication of influenza virus in the heart and prevents virus-induced cardiac fibrosis and electrical dysfunction. These results establish IFITM3 as a cardioprotective factor during influenza virus infection and provide a new model for further molecular characterization of cardiac pathogenesis of influenza.

## Results

### Generation of a novel IFITM3 KO mouse model

IFITM3 KO mice used in previous infection research are of a mixed genetic background and contain a fluorescent protein insertion at the IFITM3 locus (Lange et al., 2008). To generate a mouse model with a clean genetic deficiency on a pure C57BL/6 background, we employed a CRISPR/Cas9-based deletion strategy involving two guide RNAs targeting a region of exon 1 of the *Ifitm3* gene (Fig 1A). Mutant pups with an *Ifitm3* deletion were identified by PCR (Fig 1B). In one targeted animal, sequencing confirmed deletion of a 53 bp region of exon 1. Mutant offspring of the founder mouse were further crossed with WT mice two additional times to minimize possible off-target effects of the CRISPR/Cas9 strategy before breeding to obtain homozygous mice (Fig 1B). As a specificity control, RT-PCR for full length IFITM1, 2, and 3 coding sequences was performed on mRNA from WT and mutant (termed IFITM3 KO) MEFs. The resulting PCR products showed a band shift in the KO cells for IFITM3 only (Fig 1C) and DNA sequencing of the PCR products confirmed that IFITM1 and 2 sequences were not mutated. The *Ifitm3* mutation introduces a nonsense mutation at codon 18 and subsequent stop codon at position 37 (Fig 1A). Importantly, motifs essential for IFITM3 antiviral activity, membrane association, localization, and dimerization are not contained within its N-terminal 17 amino acids (Zani and Yount, 2018). IFITM3 protein could not be detected by immunoblotting of tissue or cell lysates from the KO mice, even upon IFNβ treatment (Fig 1D,E).

**Figure 1:**
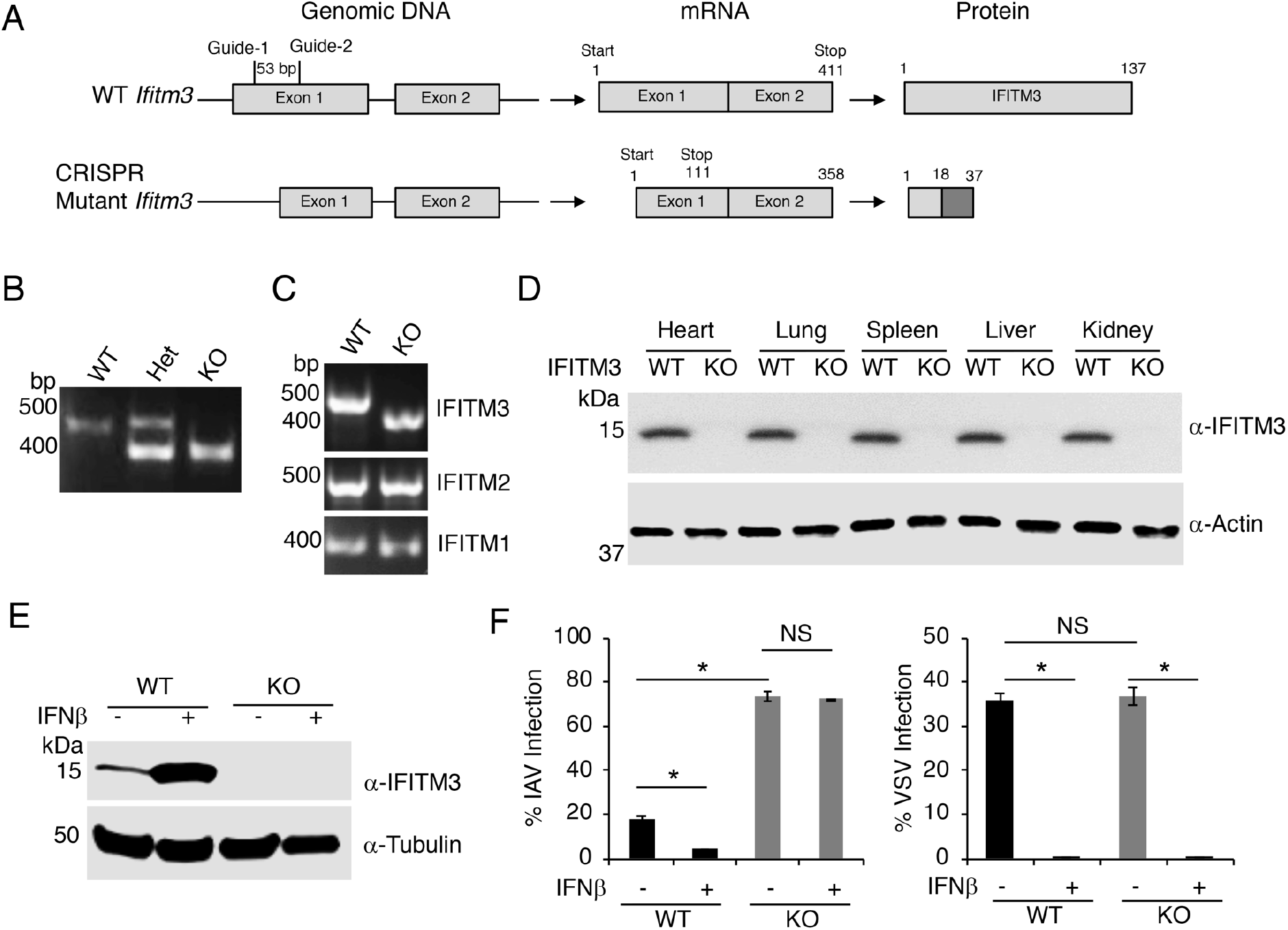
Generation and validation of a novel IFITM3 KO C57BL/6 mouse model. A) C57BL/6 zygotes were injected with a mix of Cas9 mRNA and 2 guide RNAs to target exon 1 of the *Ifitm3* gene. As a result of a 53bp deletion and consequent frame shift, the mutant Δ53 IFITM3 KO gene bears a nonsense mutation at codon 18 and a subsequent stop codon at position 37. Protein depicted for the mutant is hypothetical. B) Example genotyping PCR on genomic DNA from a WT, heterozygous (het), and KO mouse. Sequencing of the PCR products revealed a 53 bp deletion in the KO allele (not shown). C) RT-PCR for IFITMs 1-3 performed on mRNA extracted from IFNβ-treated MEFs obtained from mice of the indicated genotypes. Sequencing of the PCR products confirmed the 53bp deletion in the IFITM3 mRNA in IFITM3 KO cells and that IFITM1 and IFITM2 sequences were not mutated (not shown). D) Western blotting of tissue lysates from WT and IFITM3 KO mice. E) MEFs derived from embryos of the indicated genotypes were treated with IFNβ or were mock treated, and lysates were subjected to Western blotting. F) MEFs treated as in E were infected with influenza A virus (IAV) or vesicular stomatitis virus (VSV) for 24 h and percent infection was determined by flow cytometry. Graphs depict means of triplicate measurements from an experiment representative of at least three similar experiments. Relevant comparisons were analyzed by unpaired t-tests as indicated by lines. *p<0.001. NS, not significant.

Given the well-characterized role of IFITM3 in blocking influenza virus infection of cells (Brass et al., 2009; Yount et al., 2010), we examined the susceptibility of mouse embryonic fibroblasts (MEFs) derived from WT and KO mice to infection with H1N1 influenza A virus (PR8 strain). As expected, KO cells were significantly more susceptible to infection than WT cells (Fig 1F). This susceptibility was maintained in KO cells even after IFN treatment, which caused WT cells to be resistant to infection (Fig 1F). We observed similar trends upon influenza virus infection of WT and IFITM3 KO bone marrow-derived macrophages (Supp Fig 1). These results confirm previous experiments from our group and others indicating that IFITM3 is responsible for a significant portion of the antiinfluenza response induced by type I IFN (Brass et al., 2009; Lin et al., 2013; McMichael et al., 2017). We next examined vesicular stomatitis virus (VSV) infection of WT and KO MEFs. VSV is highly susceptible to inhibition by IFN, and although it is reported to be mildly inhibited when IFITM3 is highly overexpressed, VSV has not been tested for susceptibility to inhibition by endogenous levels of IFITM3 (Weidner et al., 2010). Unlike influenza virus infections, WT and KO cells were infected at similar rates by VSV, and VSV infection was fully restricted by IFN treatment regardless of IFITM3 status (Fig 1F). These results indicate that endogenous baseline IFITM3 does not inhibit VSV infection, and that IFITM3 is not a significant contributor to the type I IFN-induced inhibition of VSV in MEFs. These infection experiments provide important controls demonstrating that cells derived from IFITM3 KO animals are not indiscriminately susceptible to all viruses, and that the antiviral IFN response is otherwise functional in IFITM3 KO cells.

### Increased morbidity and mortality of IFITM3 KO mice upon influenza A virus infection

To examine pathogenesis of influenza A virus infection in IFITM3 KO versus WT mice, we infected mice intranasally with the PR8 strain at a dose of 10 Tissue Culture Infectious Dose 50 (TCID50), the lowest dose that allowed reproducible infections in our experiments. This dose caused weight loss in WT mice, but mice began to recover by day 12 post-infection (Fig 2A). KO mice lost significantly more weight starting at day 4 post-infection and throughout the remainder of infection (Fig 2A). By day 12 post-infection, all KO mice had succumbed to infection naturally or had lost more than 30% of their body weight and were euthanized (Fig 2B). We also measured live virus titers in the spleen (Fig 2C), and lungs (Fig 2D) on days 5, 7, and 10 post-infection. In WT mice, virus was detected in both organs, with a declining viral load seen on day 10, indicating active clearance of the virus (Fig 2C,D). In KO mice, significantly higher virus titers were measured in the spleen on days 7 and 10 as compared to WT mice (Fig 2C). Similarly, the lungs of KO mice contained higher virus titers on days 5, 7, and 10 post-infection (Fig 2D). Thus, relative to WT mice, IFITM3 KO mice are more severely infected by influenza virus.

**Figure 2:**
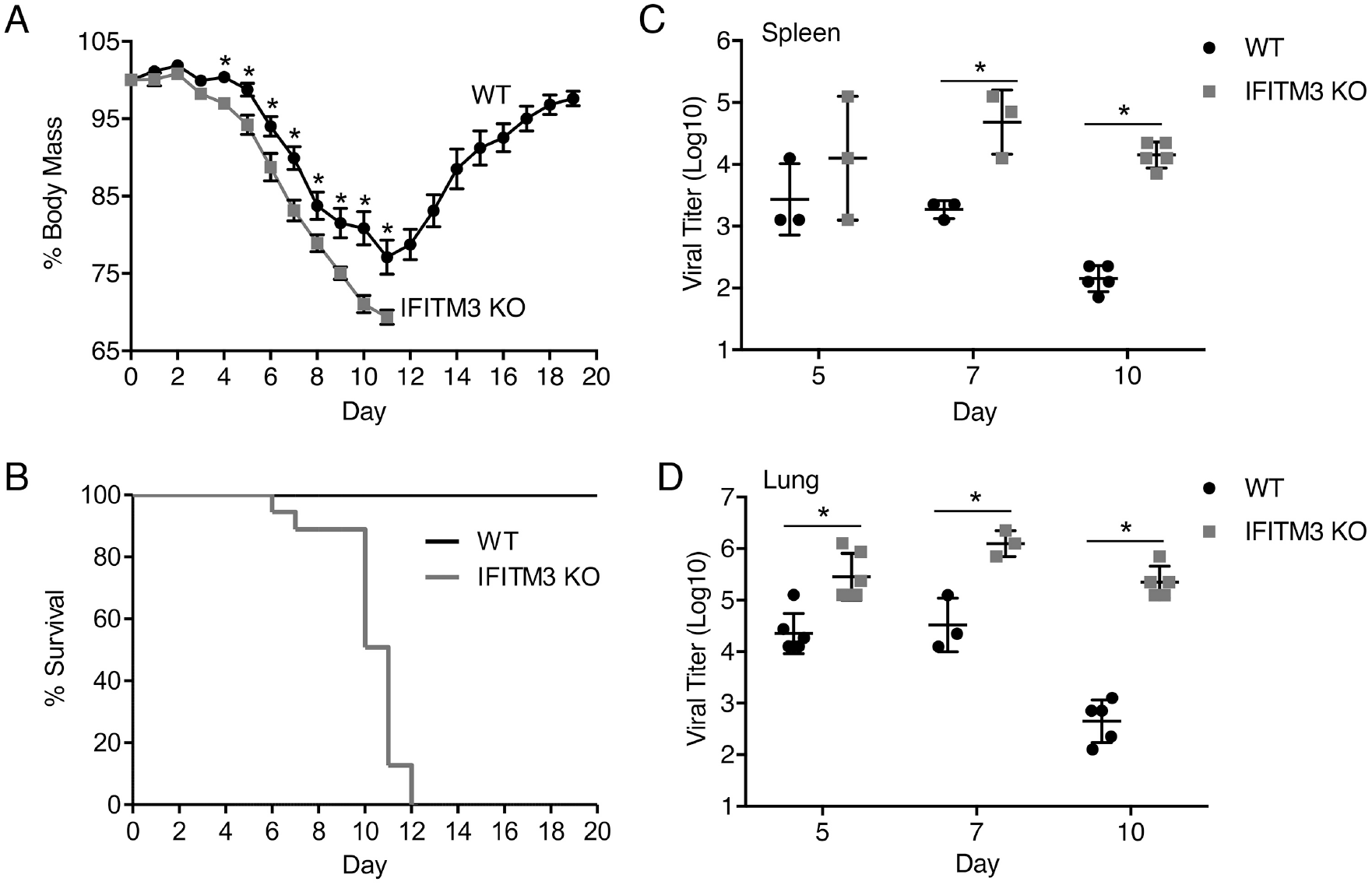
IFITM3 KO mice experience increased morbidity and mortality upon influenza virus infection. WT and IFITM3 KO mice were intranasally infected with influenza A virus strain PR8 (10 TCID50) and followed daily for A) weight loss and B) survival (n=21 KO, 29 WT). Points in A depict mean values and error bars represent standard deviation of the mean. *p<0.01 by unpaired t-test. C,D) Infected mice were sacrificed on day 5, 7 or 10 post infection for TCID50 measurement of virus titers in spleen (C) and lungs (D). Each point in C/D represents an individual mouse. Horizontal lines between points indicate mean values and error bars represent standard deviation of the mean. *p<0.001 by unpaired t test.

### Influenza virus replicates to high levels in the heart in the absence of IFITM3

We hypothesized that in addition to the spleen and lung, virus may also disseminate to higher levels in the heart. We thus quantified virus titers in the hearts of infected animals on days 5, 7, and 10 post infection. We observed a modest increase in virus titers in the hearts of KO animals as compared to WT on day 5 post infection, but observed markedly higher titers in the KO hearts on days 7 and 10 (Fig 3A). Overall, the magnitude of viral titers was highest in the lungs, followed by spleens and then hearts, which may indicate an order of dissemination. As seen in the spleen and lung, the virus titers in hearts of KO mice remained high on day 10, while the virus had declined substantially in WT mice at this timepoint (Fig 2D,C, 3A). In fact, the low virus level in the hearts of WT mice on days 5 and 7 was cleared to below the limit of detection by day 10, but remained at a titer similar to that seen on day 7 in KO mice (Fig 3A). These differences between WT and KO mice show that virus replication is inadequately controlled in each of the IFITM3 KO organs that were examined, including the heart.

**Figure 3:**
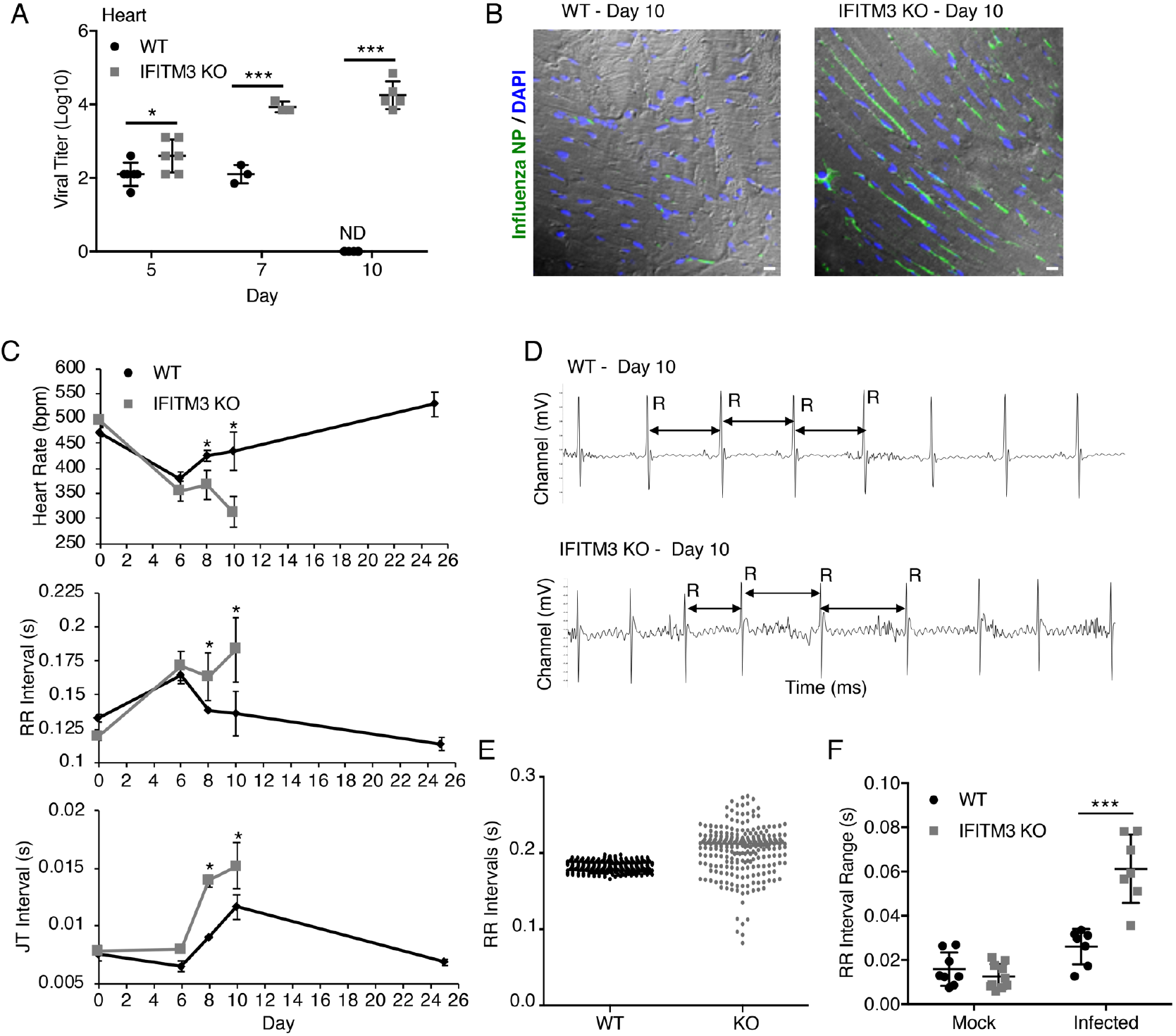
IFITM3 KO mice show uncontrolled influenza virus replication in the heart and severe cardiac electrical dysfunction during infection. WT and IFITM3 KO mice were intranasal infected with influenza A virus strain PR8 (10 TCID50). A) Infected mice were sacrificed on day 5, 7 or 10 post infection for TCID50 measurement of virus titers in the heart. Each point represents an individual mouse. Horizontal lines between points indicate mean values and error bars represent standard deviation of the mean. *p<0.05, ***p<0.0001 by unpaired t test. ND, not detected. B) Representative images of heart sections from mice sacrificed on day 10 post infection. Green, anti-influenza virus nucleoprotein (NP) staining; Blue, DAPI; Grey, brightfield imaging. Scale bar, 10 μm. C) ECG measurements over the timecourse of infection are shown (n>10). Each point depicts average values and error bars represent standard deviation of the mean. *p<0.05 by unpaired t test. D) Example ECG readings from a WT and KO mouse on day 10 post infection. Selected RR intervals are highlighted by double arrows. E) RR interval lengths are plotted for one representative example WT and one IFITM3 KO mouse on day 10 post infection from a 30 s ECG measurement period. F) RR ranges defined as the difference between the longest and shortest RR intervals over an ECG measurement period of 5 minutes was calculated for individual mice on day 10 post infection. Each point represents an individual mouse. Horizontal lines between points indicate mean values and error bars represent standard deviation of the mean. ***p<0.0001 by unpaired t test

To further identify that cells in the heart were directly infected with influenza virus, we stained heart sections from WT and KO mice on day 10 post infection with anti-influenza virus nucleoprotein antibody. We observed minimal staining of the WT hearts, consistent with clearance of the virus as indicated by our virus titration assays (Fig 3A,B). In hearts of KO mice, we primarily observed staining of elongated cells found between cardiomyocytes that are consistent with a cardiac fibroblast morphology and localization (Fig 3B). Overall, this staining allowed the direct visualization of infected cells in the hearts of KO animals.

### Influenza virus infection causes cardiac electrical dysfunction in IFITM3 KO mice

Given that replication of influenza virus to high titers has been rarely seen in the heart in mouse models (Fislova et al., 2009; Kotaka et al., 1990), we sought to examine the effect of the high virus loads in the hearts of IFITM3 KO animals on cardiac electrical function. First, we determined that IFITM3 KO mice show normal cardiac function and activity as measured by echocardiogram and electrocardiogram (ECG) (Supp Fig 2). Next, we performed ECG measurements on WT and KO mice after infection. Both WT and KO mice showed depressed heart rates and increased RR intervals on average at day 6 post infection (Fig 3C). WT mice returned to a normal heart rate and RR interval duration by day 10, while the electrical activity of the KO mice continued to significantly decline (Fig 3C). Importantly, these differing electrical measurements correlate with high virus titers detected in the KO hearts and virus clearance in the WT hearts (Fig 3A). Other ECG parameters, such as JT interval, changed later in infection, and overall showed a more dramatic electrical conduction defect in KO versus WT hearts (Fig 3C). In addition to a higher RR interval average value for the KO mice, we observed highly irregular RR intervals in KO mice with no discernible pattern, while intervals in WT mice remained regularly paced throughout infection (Fig 3D). Quantification of the RR interval range over a 5-minute ECG reading in multiple mice confirmed a consistent and significant arrhythmia in the KO mice (Fig 3E,F). Overall, we observed that high virus loads in the hearts of IFITM3 KO mice are associated with cardiac electrical dysfunction.

### IFITM3 protects mice from excess collagen deposition in the heart during infection

Cardiac fibrosis results from excess fibroblast secretion of extracellular matrix components, particularly collagen, and unresolved collagen accumulation has been shown to reduce efficiency of electrical conductivity of the heart (Maanja et al., 2017; Segura et al., 2014). Thus, we hypothesized that fibrotic collagen deposition may be linked to the cardiac electrical defects observed in infected IFITM3 KO mice. We examined heart sections from WT and KO mice at day 10 post infection after Masson’s trichrome staining, which allows visualization of collagen via blue staining. We observed significantly more blue staining in the hearts of knockout animals, indicating an enhanced fibrotic response (Fig 4A). When quantifying the blue staining per heart section on samples from multiple mice, we further confirmed that this increase in collagen staining in KO hearts was consistent for each individual mouse heart (Fig 4B). We also confirmed via qRT-PCR analysis that collagen *(Col1A2)* gene expression was increased in infected KO hearts at this timepoint (Fig 4C). Since collagen upregulation is often associated with induction by TGFβ signaling, we also examined *Tgfb1* gene expression in the hearts of infected animals and saw that it was also upregulated in KO hearts as compared to WT hearts (Fig 4D). Further, we measured IL-6 levels in heart tissue after infection for the following reasons: 1) we observed infection of cardiac cells via microscopy analysis (Fig 3B), and IL-6 is induced directly by influenza virus infection via activation of the RIG-I signaling pathway (Pichlmair et al., 2006), and 2) IL-6 is associated with cardiac fibrosis via TGFβ induction in cardiac fibroblasts (Ma et al., 2012). Indeed, we detected significantly higher amounts of IL-6 protein as measured by ELISA in heart homogenates from infected KO versus WT mice (Fig 4D). In sum, these data show that fibrotic pathways are activated at higher levels in the hearts of infected KO mice, resulting in excess collagen deposition. Together with results showing that virus replication and cardiac dysfunction are increased in KO mice, our experiments showing an enhanced fibrotic response further highlight the critical cardiac protection afforded by IFITM3 during influenza virus infection, thus identifying a new essential role for this innate immunity protein.

**Figure 4:**
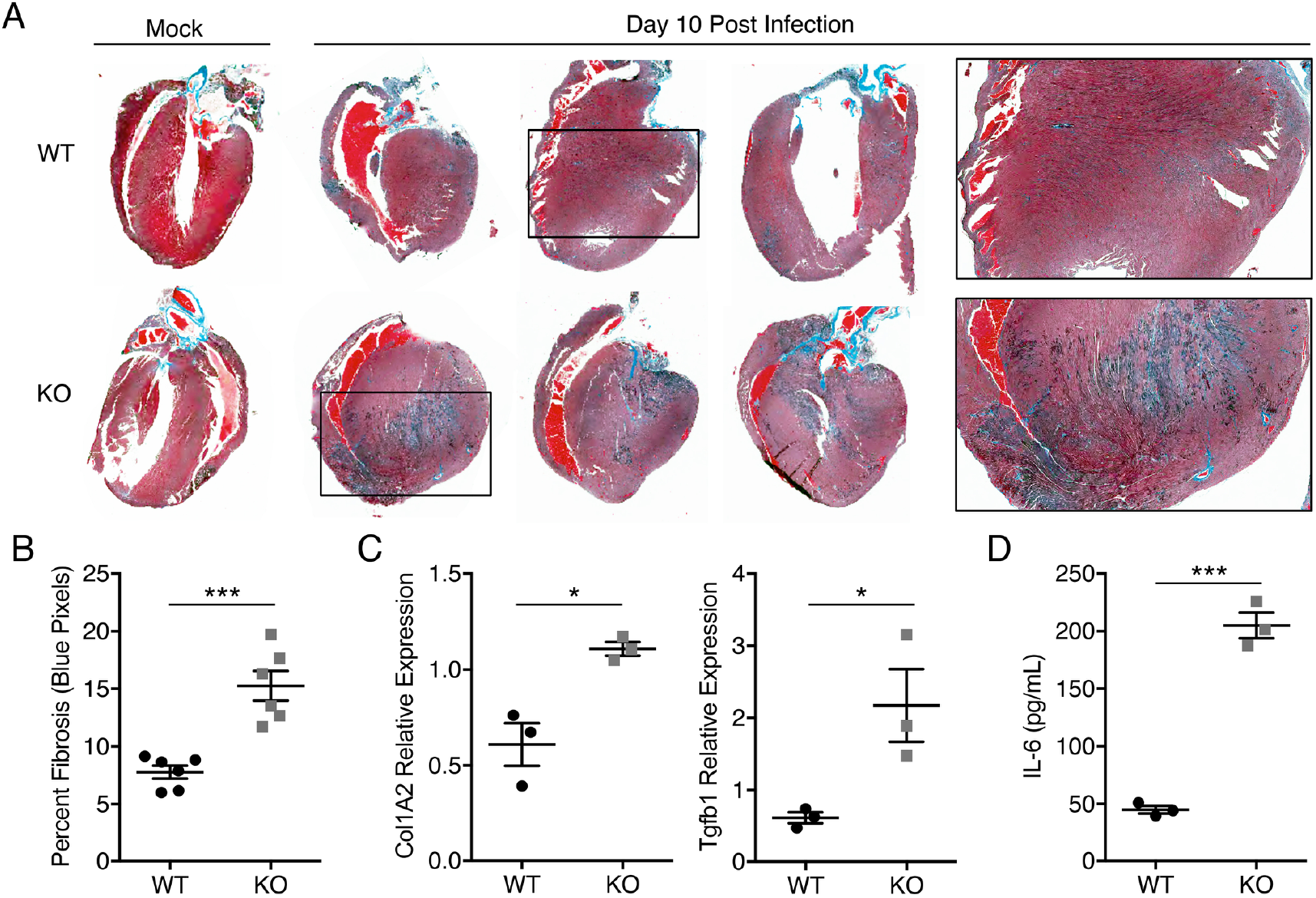
Cardiac fibrosis in influenza virus infection of IFITM3 KO mice. WT and IFITM3 KO mice were infected with influenza virus strain PR8 (10 TCID50) or were mock infected. A) Hearts were collected on day 10 post infection and sections were stained with Masson’s trichrome stain in which blue staining is indicative of fibrotic collagen deposition. Three infected hearts are shown and one representative mock infected sample is shown for each genotype. Boxed areas are regions magnified in the far right images. B) Blue pixel percentages from images as in A were quantified for infected WT and KO hearts using ImageJ software. C) Quantitative RT-PCR on mRNA extracted from hearts collected at day 10 post infection was performed to assess expression of Col1A2 and Tgfb1. D) IL-6 ELISA was performed on heart homogenates collected at day 10 post infection. In B-D, each point represents a heart from an individual mouse, horizontal lines between points indicate mean values, and error bars represent standard deviation of the mean. *p<0.05, ***p<0.0001 by unpaired t test.

## DISCUSSION

There is an underappreciated, but well-documented, effect of severe influenza virus infections in causing cardiac pathologies even in individuals without underlying cardiovascular disease (Estabragh and Mamas, 2013; Karjalainen et al., 1980; Kodama, 2010; Lucke et al., 1919; Oseasohn et al., 1959; Paddock et al., 2012; Sellers et al., 2017; Ukimura et al., 2010; Ukimura et al., 2013). Whether these effects are caused by systemic inflammation or direct infection of heart tissue remains an area of debate, though virus has been detected in human heart samples (Bowles et al., 2003; Cioc and Nuovo, 2002; Ray et al., 1989; Witzleb et al., 1976). Given that genetic defects in IFITM3 are associated with severe influenza in humans (Allen et al., 2017; Everitt et al., 2012; Kenney et al., 2017; Pan et al., 2017; Wang et al., 2014; Zani and Yount, 2018; Zhang et al., 2013), we investigated whether cardiac complications of influenza virus infection occur in an IFITM3 KO mouse model. We discovered that IFITM3 limits levels of virus in the heart (Fig 3) and prevents induction of fibrosis and resulting cardiac electrical dysfunction (Fig 3,4). Thus, in addition to identifying a novel function for IFITM3 in protecting the heart, we have also established IFITM3 KO mice as a model system for the study of cardiac complications of influenza virus infections. This mouse model provides an unprecedented opportunity to study molecular mechanisms underlying cardiac pathology induced during influenza virus infection.

Cardiac complications of infection are likely determined by a combination of host susceptibility and viral pathogenicity. Human IFITM3 polymorphisms have been linked to hospitalization or death during infection with various influenza virus strains in numerous studies (Allen et al., 2017; Everitt et al., 2012; Kenney et al., 2017; Pan et al., 2017; Wang et al., 2014; Zani and Yount, 2018; Zhang et al., 2013). Drawing parallels to our mouse experiments in which IFITM3 KO mice develop severe infections, including cardiac dysfunction, we speculate that individuals with IFITM3 defects may be more susceptible to influenza virus replication in the heart, and studies specifically addressing this question are warranted for future investigation. Additional questions remain regarding cardiac pathogenesis of influenza virus infection. For example, molecular pathways activated by cardiac infection that lead to cardiac dysfunction remain to be fully elucidated, and may involve upregulation of cytokines, such as IL-6 and TGFβ, as well as collagen downstream of these cytokines. Similarly, the role of immune cells in cardiac dysfunction caused by influenza virus infection is not known. In this regard, it is not clear why virus levels in the lungs, spleen, and heart are not controlled by the adaptive immune system in IFITM3 KO mice, though this may be explained by previous studies showing that some T cell populations may be protected from direct infection and death by their expression of IFITM3 (Allen et al., 2017; Wakim et al., 2013). Overall, IFITM3 KO mice provide a new and powerful tool for probing these intriguing questions in future experiments.

## MATERIALS AND METHODS

### Generation of IFITM3 KO mice

IFITM3 KO mice were generated by CRISPR/Cas9 technology at the Genetically Engineered Mouse Modeling Core of the Ohio State University. Briefly, single guide (sg) RNA sequences targeting the mouse *Ifitm3* gene were designed using the Benchling CRISPR/Cas9 design tool. Two sgRNAs with a predicted “On-target” score above 50 and an “Off-target” score above 60 were selected to mutate exon1. The chosen guide RNA sequences with PAM in parentheses are as follows: TTGATTCTTTCGTAGTTTGG(GGG) and GATGGGGGCACCGCACGGAT(CGG). Mouse C57BL/6 zygotes (Taconic) were injected with a mix of Cas9 mRNA (Trilink Biotechnologies) (100 ng/uL) and the two guide RNAs (50 ng/uL each) (Life Technologies). Tail-clip DNA was extracted and used to detect the presence of IFITM3 mutations by PCR and sequencing in potential founder animals. Primers used for genotyping amplify a 430 bp region and are as follows: Forward, AAACCGAAACTGCCGCAGAA; Reverse, AAAGGACCCCACACTCATACC. To minimize the potential presence of off-target mutations, IFITM3 mutants were backcrossed to C57BL/6 wild type animals a total of three generations before breeding of homozygous animals used for described experiments.

### Virus propagation and titering

Influenza A virus A/PR/8/34 (H1N1, PR8) was propagated in 10-day-old embryonated chicken eggs (Charles River Laboratories) for 48 hours at 37°C and titered in MDCK cells. VSV expressing GFP (provided by Dr. Dominque Garcin, Universite de Geneve) was propagated and titered in HeLa cells. For determining organ titers, tissues were collected and homogenized in 500ul of PBS, flash-frozen, and stored at −80°C prior to titering on MDCK cells.

### Cells, cell infections, and flow cytometry

Mouse embryonic fibroblasts (MEFs) were generated from WT and KO embryos derived from breeding of heterozygous mice and were immortalized by SV40 large T antigen transfection. MEFs and MDCK cells were grown in Dulbecco’s Modified Eagle’s Medium (DMEM) supplemented with 10% Equafetal bovine serum (FBS; Atlas Biologicals) at 37°C with 5% CO2 in a humidified incubator. Bone marrow-derived macrophages were cultured in RPMI supplemented with 10% FBS, 25 ng/mL murine M-CSF (Peptrotech), and 25 ug/mL gentamycin (Life Technologies). Where indicated, MEFs or macrophages were treated for 16 h with mouse IFNβ (eBiosciences) at 0.1 ug/mL prior to infection. Cells were infected with IAV or VSV at an MOI of 1.0 for MEFs and an MOI of 5 for BMDMs. For determination of IAV-infection percentages via flow cytometry, cells were stained with anti-H1N1 IAV NP (BEI resources) and Alexa488-conjugated secondary antibody (Life Technologies), while VSV infection rates were measured by detecting virus-encoded GFP. Flow cytometry was performed on a FACSCanto II flow cytometer (BD Biosciences) and analyzed using FlowJo software.

### Mouse infections

Mice (female, 6-10 weeks old) were anesthetized with isoflurane (Henry Schein Animal Health) and intranasally infected with influenza virus strain PR8 (10 TCID50) in 50ul of sterile saline, or were mock infected with saline in some experiments. Mice were monitored daily for weight loss and morbidity, and sacrificed if weight loss exceeded 30% of starting body mass. All procedures were approved by the OSU IACUC.

### ELISA and qRT-PCR

IL-6 concentrations in organ homogenates were analyzed using a DuoSet ELISA kit (R&D Systems). For analysis of gene expression via qRT-PCR, hearts were homogenized in TRIzol reagent (Invitrogen) and total RNA was extracted. RNA was concentrated with an RNeasy column (Qiagen) and was treated with DNAse I (Thermo Scientific). RNA was reverse transcribed using SuperScript III Reverse Transcriptase (Invitrogen). qPCR was performed using TaqMan gene expression kits (Applied Biosystems) as described previously (Makara et al., 2016).

### Western Blotting

Mouse organs, MEFs and macrophages were lysed in buffer containing 0.1 mM triethanolamine, 150 mM NaCl, and 1% SDS at pH 7.4 supplemented with EDTA-free Protease Inhibitor Cocktail (Roche) and Benzonase Nuclease (Sigma). Primary antibodies for IFITM3 (Abcam, anti-fragilis), actin (Abcam), and tubulin (Abcam) were used at 1:1000 dilutions.

### Electrocardiography

For subsurface electrocardiograph (ECG) recordings, anesthesia was provided by isoflurane in oxygen at a flow rate of 1.0 L/min, after which mice were placed in a prone position on a heated pad to maintain body temperature. For the duration of the reading, anesthesia was maintained. Subcutaneous electrodes were placed under the skin of the mouse (lead II configuration) and ECGs were recorded for at least 5 minutes on a Powerlab 4/30 (AD Instruments). ECG traces were analyzed using LabChart 7 Pro (AD Instruments).

### Echocardiography

Echocardiographic measurements were taken using a Vevo2100 imaging system (Visual Sonics) and MS-400 transducer. The mice were lightly anesthetized with isoflurane (1-1.5%) and the ejection fraction, fractional shortening, and ventricular chamber dimensions were determined in the M-mode using the parasternal short axis view at the level of the papillary muscles. Ejection fraction, fractional shortening, ventricular chamber dimensions, and LV mass were calculated using the VevoLAB program (Visual Sonics).

### Immunohistochemistry

For immunohistochemistry, hearts were fixed in 10% formalin and maintained at 4°C until embedded in parafin. Hearts were sectioned by the OSU Comparative Pathology and Mouse Phenotyping Shared Resource. Masson’s trichrome staining was used to identify fibrotic replacement of cardiac tissue. Images were analyzed via ImageJ (Version 2.0.0). Color threshold hue (scale: 0-255) was adjusted for each image to include only blue pixels (scale: 115-210) and signal intensity values were recorded. Percentage of blue pixels in the image was calculated by dividing blue pixel intensity by total pixel intensity. To ensure that color threshold parameters had been appropriately set for blue pixels, percentage of non-blue pixels was calculated by adjusting color threshold hue to exclude blue pixels (scale: 0-115 and 210-255) and dividing this value by total pixel intensity. To examine viral infection of cardiac cells, heart sections were subjected to deparaffinization and antigen retrieval prior to incubation with anti-H1N1 IAV NP (BEI resources) in a humidified chamber followed by washing and incubation with anti-mouse Alexa488-conjugated secondary antibody (Life Technologies). Nuclei were stained with DAPI. Fluorescence images were captured using an Olympus FV 1000 Spectral Confocal system.

## Acknowledgments

This research was supported by NIH grant AI130110 to J.S.Y. A.D.K. was supported by The Ohio State University Systems and Integrative Biology Training Program funded by NIH grant GM068412. T.M.M. was supported by a Gilliam Fellowship for Advanced Study from the Howard Hughes Medical Institute. A.Z. and M.C. were supported by the Ohio State University Infectious Diseases Institute training grant funded by NIH grant number AI112542 and the Ohio State University College of Medicine. The authors thank Dr. Eugene Oltz and Dr. Li Wu for editorial feedback.

## Author Contributions

Conceptualization: M.V.S.R. and J.S.Y. Methodology: A.D.K., N.M.C., Q.W., Mi.C., Fo.A., V.C., M.V.S.R., and J.S.Y. Formal Analysis: A.D.K., L.E.D., M.V.S.R., and J.S.Y. Investigation: A.D.K., T.M.M., A.I., N.M.C., L.Z., L.E.D., O.A., A.Z., Ma.C., and M.V.S.R. Resources: Fe.A., M.V.S.R., and J.S.Y. Writing – Original Draft: A.D.K. and J.S.Y. Visualization: A.D.K., M.V.S.R., and J.S.Y. Supervision: M.V.S.R. and J.S.Y. Project Administration: J.S.Y. Funding Acquisition: J.S.Y.

---

**Supplementary Figure 1:**
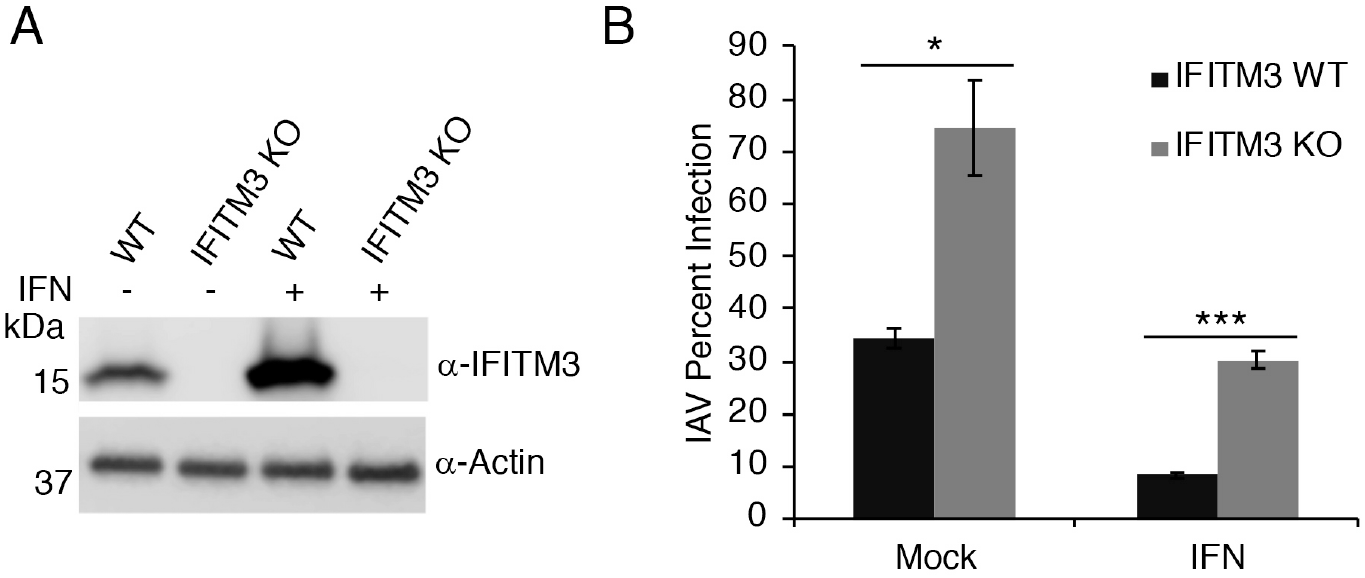
IFITM3 KO macrophages are highly susceptible to influenza virus infection. A) Bone marrow-derived macrophages of the indicated genotypes were treated with IFNβ or were mock treated, and lysates were subjected to Western blotting. B) Macrophages treated as in A were infected with influenza A virus (IAV) for 24 h and percent infection was determined by flow cytometry. Graphs depict means of triplicate samples from an experiment representative of two similar experiments. Relevant comparisons were analyzed by unpaired t-tests as indicated by lines. *p<0.01, ***p<0.0001.

**Supplementary Figure 2:**
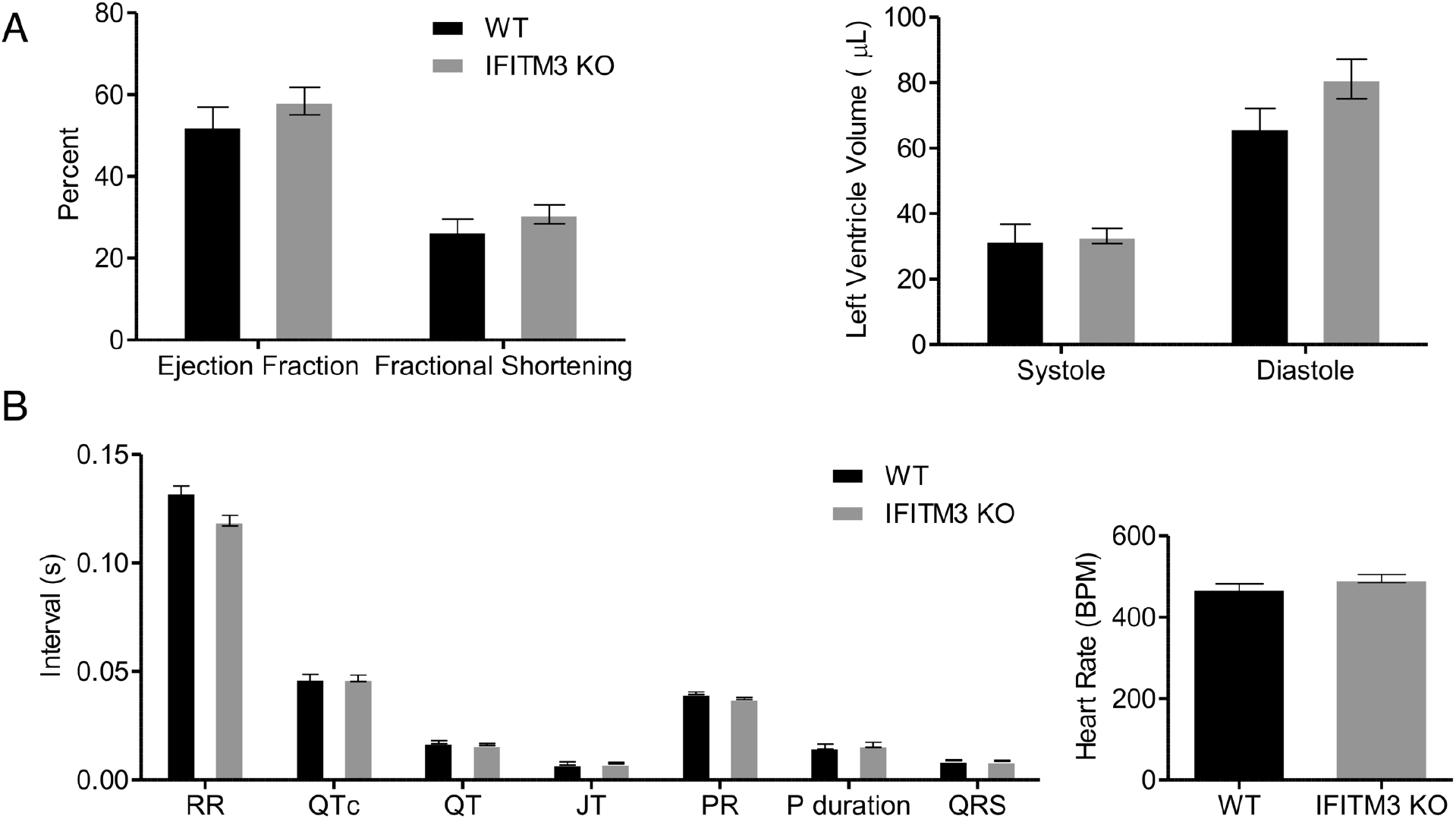
IFITM3 KO mice have normal cardiac function and electrical activity. **A**) Representative echocardiogram results from WT and IFITM3 KO mice (n=4). B) Representative electrocardiogram results from WT and IFITM3 KO mice (n=10). Error bars represent standard deviation.

